# Max is likely not at the *Drosophila* histone locus

**DOI:** 10.1101/2022.09.11.507040

**Authors:** Mellisa Xie, Skye Comstra, Casey Schmidt, Lauren Hodkinson, Leila E. Rieder

**Affiliations:** Department of Biology, Emory University, Atlanta, GA 30322

## Abstract

The histone locus body (HLB) is a conserved nuclear body that regulates histone mRNA production in metazoans. While some HLB components are known, there are likely uncharacterized factors that target the histone locus. We identified the *Drosophila melanogaster* protein Max, which interacts with known HLB member Myc, as an HLB candidate. We mapped Max ChIP-seq and ChIP-nexus datasets, which revealed encouraging signal over the histone gene array. However, we discovered that Max does not colocalize with HLB components on polytene chromosomes. Therefore, we conclude that Max is likely not at the *D. melanogaster* histone locus.

## Description

Eukaryotic cells contain nuclear bodies, which are RNA-rich membraneless organelle (Arias Escayola and Neugebauer 2018). Nuclear bodies create micro-environments to concentrate proteins and related factors that allow functions, such as RNA processing and regulation of gene expression, to occur more efficiently (Matera et al. 2009). The histone locus body (HLB) is a conserved nuclear body that is the main site of histone mRNA production in metazoans (Salzler et al. 2013). The HLB forms at the location of the clustered, repetitive histone genes. The single histone locus of the model organism *Drosophila melanogaster* includes ~100 tandem 5 kb arrays of the five replication-dependent histone genes (Duronio and Marzluff 2017; Bongartz and Schloissnig 2019). Although several HLB components of *D. melanogaster* are known, there are many uncharacterized factors that likely target the histone locus and play a role in histone gene expression. To understand how the HLB contributes to histone gene expression, our goal is to identify novel HLB components.

One method to discover novel proteins is to search for interaction partners of known HLB proteins. For example, the protein Myc is a known factor HLB factor and contributes to cell growth and proliferation (Daneshvar et al. 2011). We used the String database (Blackwood and Eisenman 1991) to identify Max, a transcription factor that interacts with Myc (Giot et al. 2003), as a candidate HLB factor. Max plays a role in cell and organismal growth (Blackwood and Eisenman 1991) and is known to form a complex with Myc, which functions as a transcriptional activator (Amati et al. 1992; Kretzner et al. 1992).

He *et al*. 2015 studied Max localization in *Drosophila melanogaster* S2 cells using ChIP-seq to compare it to their novel technique, ChIP-nexus, which maps transcription factor binding with higher resolution than ChIP-seq (He et al. 2015). We mapped their publicly available datasets (NCBI GEO accession: GSE55306) to the histone gene array (HA) (McKay et al. 2015) using Galaxy, an open-source computational biology platform (Afgan et al. 2018). We used “Bowtie2,” to map Max’s sequencing data to the HA and produced a BAM file. “BamCoverage” was run on the BAM file to produce a bigwig file, which contains better visualization of the alignment. We did not normalize the data to input, as no input data was provided by He *et al*. 2015. We visualized the resulting bigwig files using Integrative Genomics Viewer (IGV) (Robinson et al. 2011). Max ChIP-seq and ChIP-nexus data aligned to the HA gave substantial signal, notably over promoters and gene bodies (Fig. 1A). The profiles of Max ChIP-seq and ChIP-nexus did not differ significantly. These observations suggest that Max localizes to the HA in *Drosophila* S2 cells.

**Figure 1.**
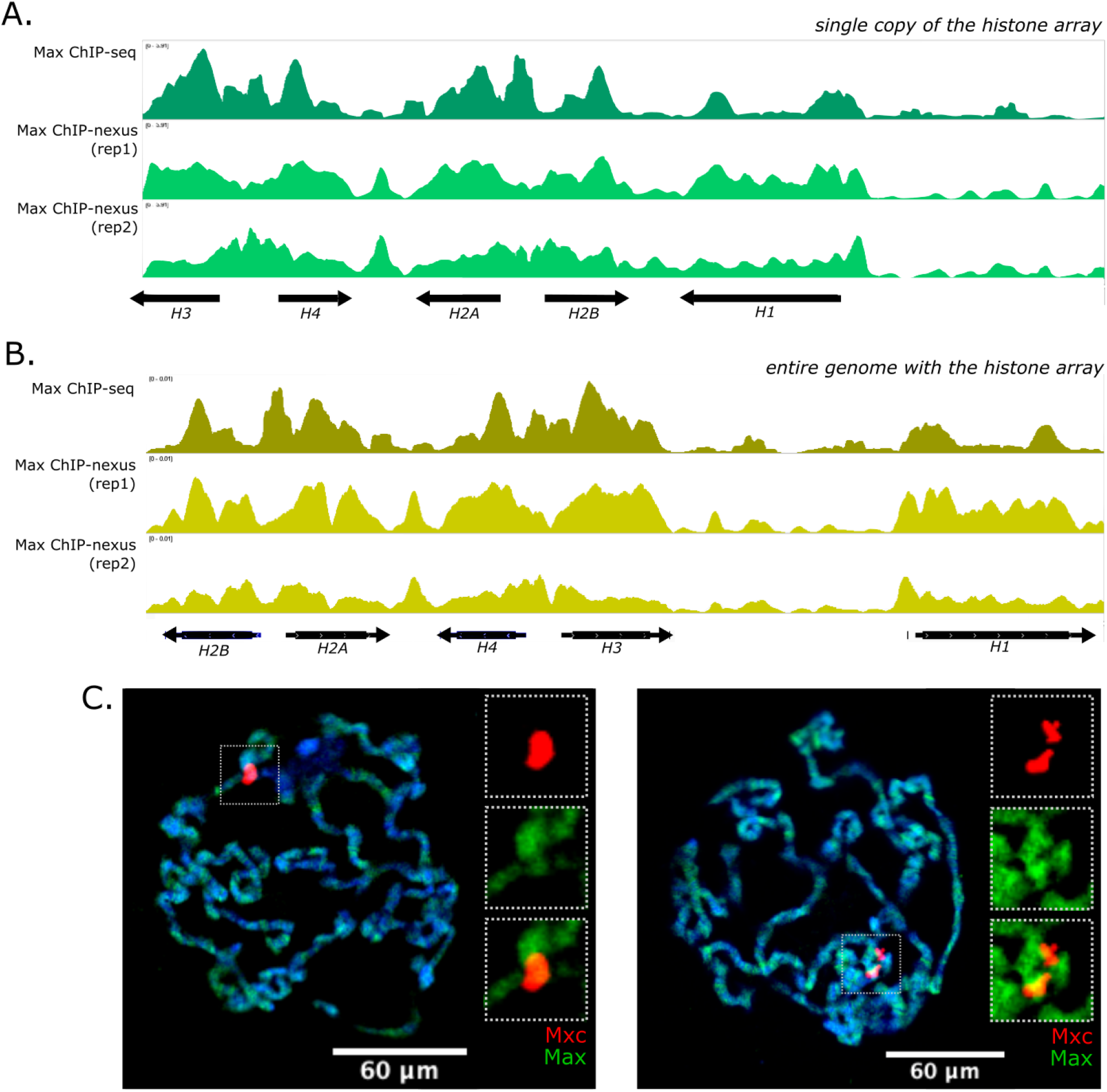
Max ChIP data and polytene chromosome immunostaining. (A) We mapped Max ChIP-seq and ChIP-nexus data from He *et. al* 2015 to a custom genome consisting of a single copy of the *Drosophila melanogaster* histone array (McKay et al. 2015). (B) We also mapped the Max ChIP-seq and ChIP-nexus data to the entire *D. melanogaster* genome. ChIP-seq and ChIP-nexus data are not input normalized, as no input dataset was provided by He *et al*. 2015. (C) Polytene chromosome immunostaining using an anti-Mxc antibody (red) reveals the histone locus body. Immunostaining with an anti-Max antibody (green) reveals that Max binds broadly across the whole genome and does not colocalize at the histone locus with Mxc. Two representative chromosome spreads are shown.

To ensure that we did not generate artifactual Max profiles by mapping whole genome data to one 5 kb HA, we mapped the same Max ChIP-seq and ChIP-nexus datasets to the entire *D. melanogaster* genome (dm3), including a single HA copy (Fig. 1B). Max ChIP-seq and ChIP-nexus profiles over the HA did not change appreciatively when mapped to the whole genome.

To confirm Max localization to the histone locus, we stained polytene chromosomes from third instar larval salivary glands. We co-stained chromosomes with anti-Mxc antibody, a marker for the HLB (White et al. 2011), and anti-Max antibody. Immunostaining reveals that Max binds broadly across polytene chromosomes and does not colocalize with Mxc at the histone locus (Fig. 1C).

Although ChIP-seq and ChIP-nexus data suggested that Max is present at the histone locus (Fig. 1A-B), polytene chromosome immunostaining did not confirm Max localization (Fig. 1C). We conclude that Max is likely not present at the histone locus but note that it may have different localization patterns in different tissues. Ultimately, although a bioinformatics approach can identify novel protein HLB candidates, further wet-lab experiments are necessary to confirm the localization of these components to the histone locus.

## Methods

The computational biology tool, Galaxy, was used to map Max to determine if there was signal at the genome of the HLB. First, Max ChIP-seq and ChIP-nexus FASTQ files were extracted from He *et al*. 2015’s Sequence Read Archive (SRA) list (NCBI GEO accession: GSE55306) in Galaxy using the “Faster Download and Extract Reads in FASTQ” option. The extracted FASTQ files contain Max’s sequence and quality information. Next, the reference genome, the histone array (HA), was normalized in Galaxy using the “NormalizeFASTA” option. This will ensure that the HA genome will be compatible with Max’s FASTQ files that were extracted. Afterward, Galaxy’s option, “Bowtie2,” was used to map Max’s sequential data to the HA and produced a BAM file. Finally, “bamCoverage” was run on the BAM file to produce a bigwig file, which contains better visualization of the alignment. The produced bigwig file was displayed in the Integrative Genomics Viewer (IGV).

Immunostaining was performed on polytene chromosomes of third-instar larval salivary glands that developed in 18°C temperatures. For the primary antibody staining, we co-stained the chromosomes using anti-Mxc antibody at a 1:5000 dilution and anti-Max antibody at a 1:500 dilution. For the anti-Mxc antibody, a secondary anti-guinea pig antibody was used at a 1:1000 dilution. For the anti-Max antibody, a secondary anti-rabbit antibody was used at a 1:1000 dilution.

## Reagents

*Drosophila melanogaster* genotype: LR44: y[1]w[1118]; +;+;+

Guinea pig anti-Mxc antibody (1:5000), a gift from Dr. Robert Duronio.

Rabbit anti-Max antibody (1:500), a gift from Dr. Julia Zeitlinger (previously commercially available from Santa Cruz: sc-28209).

AlexaFluor647 anti guinea pig (1:1000), available at Thermo Fisher Scientific (catalog #: A-21450).

AlexaFluor488 anti-rabbit (1:1000), available at Thermo Fisher Scientific (catalog #: A-11008).

## Acknowledgments

We thank John Ali for generating the Galaxy pipeline. We thank Dr. Julia Zeitlinger, who provided the anti-Max antibody (sc-28209), and Dr. Bob Duronio for the anti-Mxc antibody. We thank Gwyn Puckett, Tommy O’Haren, Greg Kimmerer, Mary Wang, and Dabin Cho, who supported this project. We thank John Ali for generating the Galaxy pipeline. We thank Dr. Julia Zeitlinger, who provided the anti-Max antibody (sc-28209), and Dr. Bob Duronio for the anti-Mxc antibody. We thank Gwyn Puckett, Tommy O’Haren, Greg Kimmerer, Mary Wang, and Dabin Cho, who supported this project.

## Author Contributions

Mellisa Xie: conceptualization, formal analysis, funding acquisition, investigation, visualization, writing - original draft. Skye Comstra: conceptualization, funding acquisition, methodology, writing - review editing. Casey Schmidt: conceptualization, writing - review editing. Lauren Hodkinson: conceptualization, writing - review editing. Leila E. Rieder: conceptualization, funding acquisition, project, writing - review editing.

